# Gene expression analysis of *Cyanophora paradoxa* reveals conserved abiotic stress responses between basal algae and flowering plants

**DOI:** 10.1101/674762

**Authors:** Camilla Ferrari, Marek Mutwil

## Abstract

- The glaucophyte *Cyanophora paradoxa* represents the most basal member of the Archaeplastida kingdom, however the function and expression of most of its genes are unknown. This information is needed to uncover how functional gene modules, i.e. groups of genes performing a given function, evolved in the plant kingdom.
- We have generated a gene expression atlas capturing responses of Cyanophora to various abiotic stresses. This data was included in the CoNekT-Plants database, enabling comparative transcriptomic analyses across two algae and six land plants.
- We demonstrate how the database can be used to study gene expression, co-expression networks and gene function in Cyanophora, and how conserved transcriptional programs can be identified. We identified gene modules involved in phycobilisome biosynthesis, response to high light and cell division. While we observed no correlation between the number of differentially expressed genes and the impact on growth of Cyanophora, we found that the response to stress involves a conserved, kingdom-wide transcriptional reprogramming, which is activated upon most stresses in algae and land plants.
- The Cyanophora stress gene expression atlas and the tools found in https://conekt.plant.tools/ database provide a useful resource to reveal functionally related genes and stress responses in the plant kingdom.

## Introduction

The glaucophyte algae are a basally diverging group of unicellular taxa with four described genera and an estimated 15 species (Kies & Kremer, 2006). Glaucophytes represent one branch of the Archaeplastida (Plantae) kingdom, where the other branches are represented by the red (Rhodophyta) and green algae and plants (Viridiplantae) (Adl *et al.*, 2012). The common ancestor of these taxa captured the cyanobacterial endosymbiont ca. 1.6 billion years ago, giving rise to the chloroplast found in glaucophytes, red and green algae, and land plants (Margulis Lynn, 1981; Bhattacharya *et al.*, 2004; Yoon *et al.*, 2004; Blank, 2013). The plastids of red and green algae participated in several other endosymbioses, leading to the appearance of diatoms, dinoflagellates, euglenids, and haptophytes (Reyes-Prieto *et al.*, 2007). Glaucophytes, and their representative *Cyanophora paradoxa*, retain traits from the ancestral cyanobacterial endosymbiont, such as phycobilisomes, peptidoglycan (PG), an ancient, primitive RNA interference pathway (Gross *et al.*, 2013), lack of chlorophyll-b (Löffelhardt, 2014) and presence of a bacterial-derived UhpC-type hexose-phosphate transporter used to translocate sugars from the plastid to the host cytosol, that is not found in land plants (Price *et al.*, 2012). Therefore, due to these unique traits, Cyanophora can provide invaluable insights into the ancestral state of the Archaeplastida host, its photosynthetic organelle, and the evolution of the functional gene modules found in the plant kingdom.

Our ability to extract useful knowledge from the Cyanophora genome, and to understand how its genes work together to form these traits, relies on our ability to correctly assign biological functions to gene products (Rhee & Mutwil, 2014; Hansen *et al.*, 2018). Whereas the experimental characterization of protein function is resource□intensive, the *in silico* inference of gene function through gene function prediction methods can be fast and readily available. Therefore, with the increasing availability of assembled genomes, gene function prediction is currently one of the most active areas in bioinformatics. Of the many available methods, co-expression analysis is the most successful and reliable way of predicting gene function (Lee *et al.*, 2010; Hansen *et al.*, 2014, 2018; Rhee & Mutwil, 2014; Proost & Mutwil, 2016; Ruprecht *et al.*, 2017b). Co-expression is based on the guilt-by-association principle, which states that genes involved in the same or closely related biological processes tend to have similar expression patterns across organs, developmental stages and biotic as well as abiotic perturbations (Usadel *et al.*, 2009). Co-expression analyses are widely used (Radivojac *et al.*, 2013; Rhee & Mutwil, 2014; Jiang *et al.*, 2016), and have been applied to successfully identify genes involved in plant viability (Mutwil *et al.*, 2010), seed germination (Bassel *et al.*, 2011), shade avoidance (Jiménez-Gómez *et al.*, 2010), cyclic electron flow (Takabayashi *et al.*, 2009), cell division (Takahashi *et al.*, 2008), drought sensitivity and lateral root development (Lee *et al.*, 2010), and others (Stuart *et al.*, 2003; Yu *et al.*, 2003; Persson *et al.*, 2005; Itkin *et al.*, 2013; Proost & Mutwil, 2016; Sibout *et al.*, 2017).

Classical comparative genomic approaches that are based on gene sequences are very useful, however they present certain shortcomings as genes often operate as functional gene modules (Hartwell *et al.*, 1999). This means that gene products frequently form e.g. enzymatic pathways or protein complexes that usually require multiple genes working together to perform a given task (e.g., photosystem II complex and ribosomes). While genomic analyses can reveal which gene families have appeared and expanded in a given organism, it might not show which of the underlying genes are functionally related. Since plant gene families can be large and functionally divergent (Shiu & Bleecker, 2001; Shiu *et al.*, 2005), sequence□based analyses could result in incorrectly predicted gene functions (Lynch & Katju, 2004). Therefore, a more rewarding approach to study the evolution of new traits needs to integrate the classical genomic approaches with functional gene modules, which can be identified by expression and co-expression analysis (Ruprecht *et al.*, 2017b; Lampugnani *et al.*, 2019).

To establish these enhanced, co-expression-driven comparative analyses for Cyanophora, we first investigated the changes in gene expression as a response to 10 different abiotic stresses. Together with publicly available data that capture changes in gene expression during 13 time points of the diurnal cycle, we provide a comprehensive expression atlas of this basally diverging member of the Archaeplastida kingdom. To make the data easily accessible we have uploaded it to the CoNekT-plants database, which provides advanced comparative transcriptomic analyses for algae and land plants. We then exemplify how these analyses can be used to study phycobilisome formation, response to high light, and cell division. Finally, we perform a meta-analysis of stress responses in the Archaeplastida kingdom and find evidence of a kingdom-wide transcriptional reprogramming mechanism as a response to a wide variety of stresses.

## Materials and Methods

### Experimental growth conditions and sampling

Cyanophora paradoxa UTEX555 (SAG 29.80, CCMP329) was obtained from Provasoli-Guillard National Center for Marine Algae and Microbiota (NCMA, Bigelow Laboratory for Ocean Sciences). The cultures were grown in C medium (Table S1) at 24°C under continuous light (40 μmol photons m^−2^ s^−1^) and aeration. We have modulated the magnitude of stresses to inhibit but not stop the cell division of the cultures. The growth of the cultures, measured as cells/mL, was measured with a Z2 Coulter Counter (Beckman Coulter).

When subjected to nutrient stress (deprivation of nitrogen: 10% of recommended KNO_3_, sulphur: 2% of recommended MgSO_4_ 7H_2_O, phosphate: no GlycerophosphateNa_2_ 5H_2_O and trace metals: no trace metal solution, Table S1), the culture was washed twice with new media and the growth rate was monitored daily for six days. On the fifth day, part of the culture was harvested for RNA isolation, while the remaining part was left to monitor the growth rate. The control culture contained unmodified C medium.

When subjected to environmental stress (high light, heat, cold, prolonged darkness), the cultures were grown in C medium without aeration for seven days and subjected to the stresses for 12 hours, except for prolonged darkness, which lasted for 72 hours. The cultures treated for cold (4° for 12 hours), heat (37° for 12 hours), and the control (24°C for 12 hours), were kept in the dark for the duration of the experiment. The high light treatment (150 μmol photons m^−2^ s^−1^, 24°C for 12 hours) and its control culture (40 μmol photons m^−2^ s^−1^, 24°C for 12 hours), were performed simultaneously with the other environmental stress samples. After harvesting, the remaining culture was transferred to normal growth conditions (24°C, under continuous light, 40 μmol photons m^−2^ s^−1^, aeration with ordinary air) and the growth was monitored for the following three days. The reference time 0 was assigned to the moment the stress was first applied.

### Determination of starch levels

Cultures with ~10^6^ cells/mL were split into two cultures. One culture was incubated in the dark for 72 hours, while the control culture was in continuous light at 40 μmol m^−2^ s^−1^. For visualizing the starch levels, the cells were stained with 20% Lugol solution and analysed by microscopy (Fig. 1c). For the optical density (OD) measurements, 20 mL of culture were harvested (300 g for 10 minutes at 4°C) and the cell pellet was incubated with 200 μL of DMSO at 70°C for 5 minutes, followed by incubation with 3 mL of MetOH at 4°C for 12 hours. The cells were collected by centrifugation and resuspended with 200 μL of 70% EtOH, 400 μL 2N NaOH and 400 μL H_2_O and incubated at 25°C for 2.5 hours. A solution of 400 μL 2N HCl, 1 mL 0.5 M NaAcetate, 7 mL H_2_O and 200 μL iodine reagent (1% KI + 0.1% I_2_) was added right before measuring the absorbance at 680 nm (Avidan *et al.*, 2015). Three independent cultures with different cell densities were used for this analysis: i) ~0.5 * 10^6^ cells/mL, ii) ~10^6^ cells/mL and iii) ~2*10^6^ cells/mL. Three technical replicates were performed.

**Figure 1.**
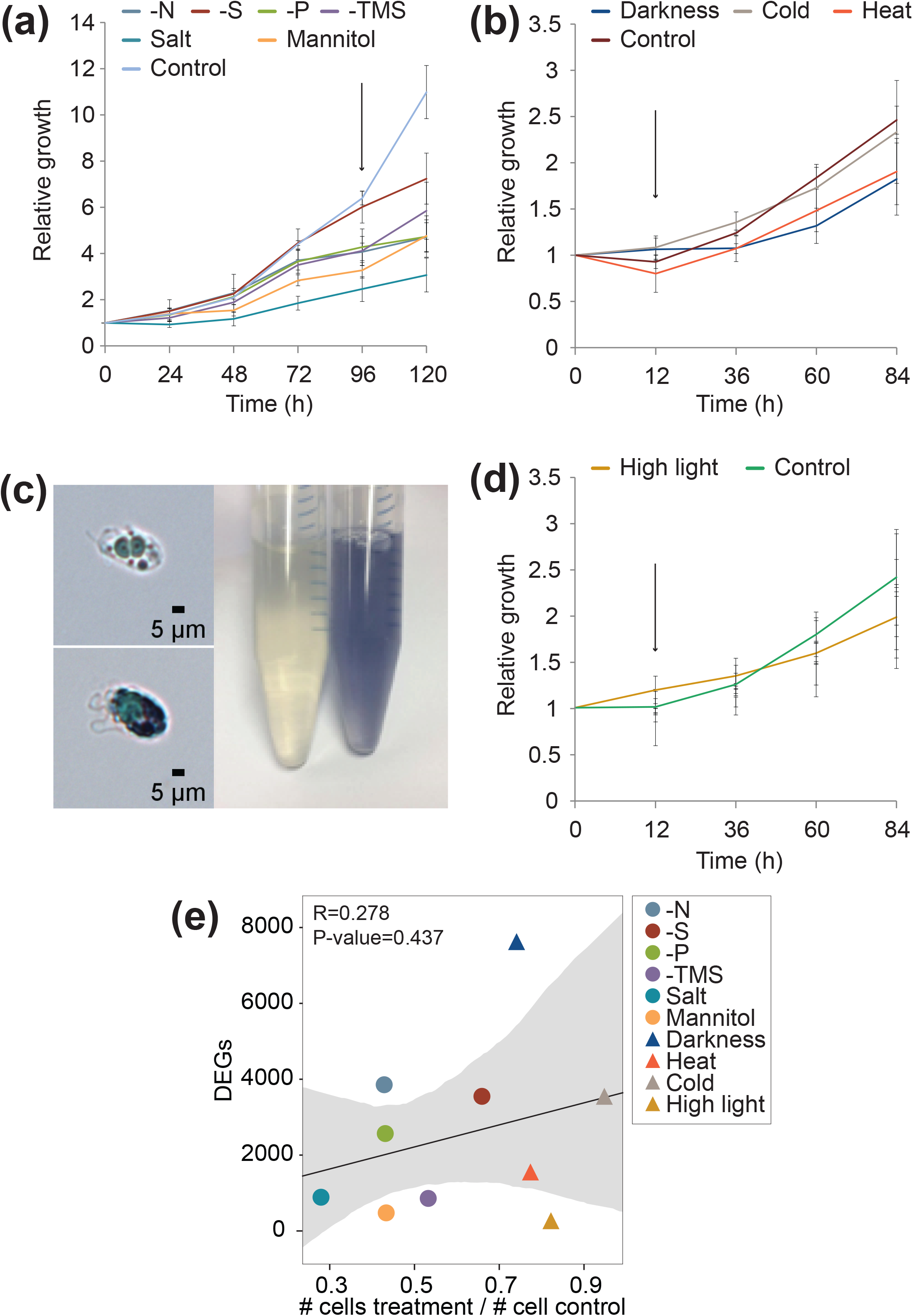
Differential gene expression analysis of the *Cyanophora paradoxa* stress expression atlas. (a) Relative growth rates of the cultures subjected to nutrient stresses. The x-axis shows the time after the cells were moved to the stress media, while the y-axis indicates the relative growth rate, given as the number of cells in a culture/number of cells in culture at time 0. Error bars indicate standard deviation, while the black arrow shows sampling time. (b) Relative growth rates in the darkness, cold and heat experiments. (c) Starch granules in the dark and control cultures. The Lugol solution stains starch granules with dark blue color. (d) Relative growth rates in the high light experiment. (e) The relationship between the relative growth rate and the number of differentially expressed genes. The gray area indicates the confidence interval of the regression.

### RNA preparation and sequencing

The RNA was extracted using Spectrum™ Plant Total RNA Kit (Sigma-Aldrich) according to the manufacturer’s instructions. The integrity of the RNA was assessed using RNA nano chip on Agilent Bioanalyzer 2100. The libraries were prepared from total RNA using polyA enrichment and sequenced using Illumina-HiSeq2500/4000 at Beijing Genomics Institute and Max Planck-Genome-centre in Cologne. Three RNA isolations were done for each sample.

### Analysis of RNAseq data and microarrays data

The reads were trimmed, mapped, counted, and TPM-normalized using the LSTrAP pipeline (Proost *et al.*, 2017). The genome used for mapping the reads is *C. paradoxa* v1.0 (Price *et al.*, 2019). More than 81% of the reads mapped to the genome, and on average 88% of the reads mapped to the coding sequences (Table S2). The principal component analysis revealed that one of the three replicates of salt stress did not cluster with the other two replicates. For this reason, we excluded the outlier sample and sequenced two additional biological replicates.

The publicly available data was processed as follows: raw expression data for *Arabidopsis thaliana*, *Chlamydomonas reinhardtii*, and *Synechocystis sp. PCC 6803* (Table S3) were downloaded from ArrayExpress. The raw fastq files were processed using the LSTrAP pipeline and the reads were mapped to *A. thaliana* TAIR10 and *C. reinhardtii* v5.5. Raw counts from the RNA-seq experiments were used with the R package DESeq2 (Love *et al.*, 2014) to identify differentially expressed genes (DEGs). The various types of microarrays were processed based on the manufacturer. Affymetrix cel files were processed with the R package affy (Gautier *et al.*, 2004). Agilent raw files were processed with the R package limma (Ritchie *et al.*, 2015). Differentially expressed genes for all microarray experiments were identified by using the R package limma. For further analyses, we only considered the genes which showed an adjusted p-value < 0.05 and a −1>log_2_ fold>1 as DEGs (Table S4-S7).

### Adding Cyanophora to the CoNekT-plants database

The normalized TPM matrix containing expression values from diurnal cycle (Ferrari et al., 2019) and the expression matrix of the newly generated stress samples (Table S8) were uploaded in CoNekT-Plants together with CDS sequences (Price *et al.*, 2019), annotation (Mercator4 v1.0) (Schwacke *et al.*, 2019) and protein domain information (InterProScan 5.32-71.0) (Jones *et al.*, 2014). The gene families were detected with OrthoFinder 1.1.8 (Emms & Kelly, 2015). For the functional description of the genes, we used Mercator4 v1.0 with standard settings. To investigate the effect of the various stresses on the main biological processes, we analyzed the expression of genes assigned to the first level MapMan bins obtained from Mercator.

### Functional enrichment analysis of clusters

The Heuristic Cluster Chiseling Algorithm (HCCA) clusters of *C. paradoxa* were downloaded from CoNekT-Plants (Table S9). For each cluster we counted the number of times a specific biological process was represented. This observed distribution of biological processes was compared to a permuted distribution obtained by sampling an equal number of genes from the total pool of *C. paradoxa* genes for 10,000 permutations. The empirical p-values obtained were false-discovery rate (FDR) corrected by the Benjamini-Hochberg procedure (Benjamini & Hochberg, 1995).

### Gene function analysis of abiotic stress

To investigate how the stress conditions perturbed certain biological processes (MapMan bins), we first analyzed the fraction of differentially expressed genes (DEGs) assigned to each bin. We assigned the fractions into strongly affected (≥50% of genes in the bin are DEGs), mildly affected (20% ≤ DEGs < 50%) or weakly affected (DEGs < 20%). To visualize if the DEGs found in the bin are preferentially up- or down-regulated we used the formula:

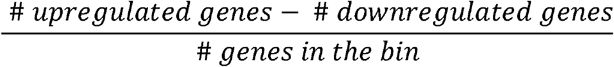

The formula returns 1 when all genes in the bin are upregulated, 0 when there is an equal number of up- and down-regulated genes and −1 when all genes in the bin are downregulated.

### Inferring similarities of stress responses

We used the Jaccard Index (JI) to assess the similarity of stress responses between two experiments. To this end, we first identified the gene families corresponding to the DEGs, which were used to calculate the Jaccard index for two compared stresses, X and Y. The observed JI between stresses X and Y is:

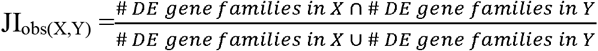

To test whether a given JI_obs(X,Y)_ is larger than expected by chance, i.e. two stresses show similar transcriptomic response, we performed a permutation test where we drew a number of genes equal to the amount of DEGs present in a specific experiment (DEGs_r) and calculated the expected Jaccard indexes for each experiment pair (JI_perm(X,Y)_). The procedure was performed 10,000 times, and the empirical p-values obtained, calculated independently for up- and downregulated families, were FDR corrected. The OrthoFinder gene families used in this analysis were obtained from (Ferrari *et al.*, 2019).

### Identification of stress-responsive gene families

We counted the number of experiments when a given family contained DEGs, and calculated the empirical p-value by estimating if this number was larger than the corresponding counts obtained from shuffled gene-DEG assignments. The shuffling was done 10,000 times, and the empirical p-values were FDR corrected.

### Phylostratigraphic enrichment analysis of DEGs

To test whether there is a correlation between the age (phylostrata) (Domazet-Loso *et al.*, 2007) of a gene family and the transcriptomic responses to stress, we calculated the fraction of DEGs found in each phylostratum. Every fraction was then tested for significance by estimating whether the observed fraction was larger than fractions of shuffled gene-phylostrata assignments. The shuffling was done 10,000 times, and the empirical p-values were FDR corrected. The gene-phylostrata assignment was retrieved from (Ferrari *et al.*, 2019).

### Data availability

The raw sequencing data is available from EBI accession number E-MTAB-7822.

## Results

### Generation of *C. paradoxa* expression atlas

To be able to assign gene functions via co-expression analyses we induced broad changes in gene expression in *C. paradoxa* by subjecting the liquid culture to four nutritional, one osmotic, one salt and four environmental stresses. The intensities of the different stresses were modulated to reduce the growth rate without stopping it entirely. To this end, culture growth was monitored by cell counting from the time when the stress was first applied (time 0 hours) up to several days after the harvest time for gene expression analyses (Fig. 1a, b, and d). The nutritional stress category involves the depletion of macro or micronutrient to the growth medium (nitrogen, sulphate, phosphate, micronutrient solution), while the salt and osmotic stresses were induced by addition of 75 mM NaCl or 100 mM mannitol, respectively. Harvest time for all nutritional, salt and osmotic stress conditions was on day 4 when a pronounced reduction of growth was observed for most stresses compared to the control (Fig. 1a). The presence of NaCl in the medium had the strongest negative effect on growth, followed by presence of mannitol, depletion of phosphate and depletion of TMS. The effect of sulphate depletion was only visible on day 5 (Fig. 1a). For the environmental stresses we subjected the cultures to heat (37°C), cold (4°C), high light (150 μE) with a harvest time at 12 h for each stress condition (Fig. 1b and 1d). Cultures grown for 72 hours in darkness showed an expected absence of starch granules as compared to the control (Fig. 1c, Fig. S1). The environmental stresses resulted in less pronounced negative growth effects (Fig. 1b, d), however on day 2.5 and day 3.5 after the environmental stress was applied, the effects were more clearly visible at the level of growth rate (Fig. 1b, d).

To evaluate how gene expression in Cyanophora was affected by the different stresses, RNA was isolated from the harvested cultures and sequenced in triplicates to produce at least 15 million reads per sample (Table S2). Principal component analysis (PCA) revealed an expected clustering of replicates and related stresses (Fig. S2). While the separation of the different treatments was quite distinct for most stresses, we did not observe a clear separation between the high light treatment and its specific control (Fig. S2b), suggesting a milder effect of increasing light intensity to 150 μE.

To identify the significantly differentially expressed genes (DEGs, adjusted p-value < 0.05, −1>log_2_ fold>1), we compared the raw counts of the stress conditions to the respective control. The impact of the stress on the transcriptome was highly variable, ranging from 270 DEGs detected after 12 hours of high light treatment to 7628 DEGs after 72 hours of darkness (Table S5). Surprisingly, we did not observe any significant correlation between the number of DEGs and the magnitude of effect the stress had on the relative growth of the culture (Fig. 1e). Cold and sulphate deprivation (gray triangle and red dot, respectively) showed mild inhibition of growth, but a much higher proportion of DEGs compared to treatments with a severe growth phenotype, such as salt and mannitol treatments (Fig. 1e, turquoise and orange dots respectively). Consequently, we observed no significant correlation between DEGs and relative growth rate in Cyanophora (r = 0.278, p-value = 0.437).

### Integration of Cyanophora transcriptome into CoNekT database

Biological networks are characterized by their scale-free topology nature, which results in few genes being connected (correlated) to many genes, while the majority of genes have only a few connections (Barabási & Bonabeau, 2003). This architecture is presumed to ensure stability in the case of perturbation, as the network topology remains unaffected when certain genes mutate (Barabási & Oltvai, 2004). To test if the produced *C. paradoxa* data also follows an expected scale-free network, we calculated the Pearson Correlation Coefficient (PCC) for every gene pair, with a threshold of 0.9 (Usadel *et al.*, 2009), and counted the number of times a given gene is co-expressed with other genes at this threshold (node degree). The points of the resulting power law plot formed a line with a negative slope, showing a negative correlation between node frequency (i.e., number of genes with a certain number of connections) and node degree (i.e., number of connections per gene). Thus, this result confirms the scale-free topology of the network and indicates that our expression data produces biologically relevant topology (Fig. 2a).

**Figure 2.**
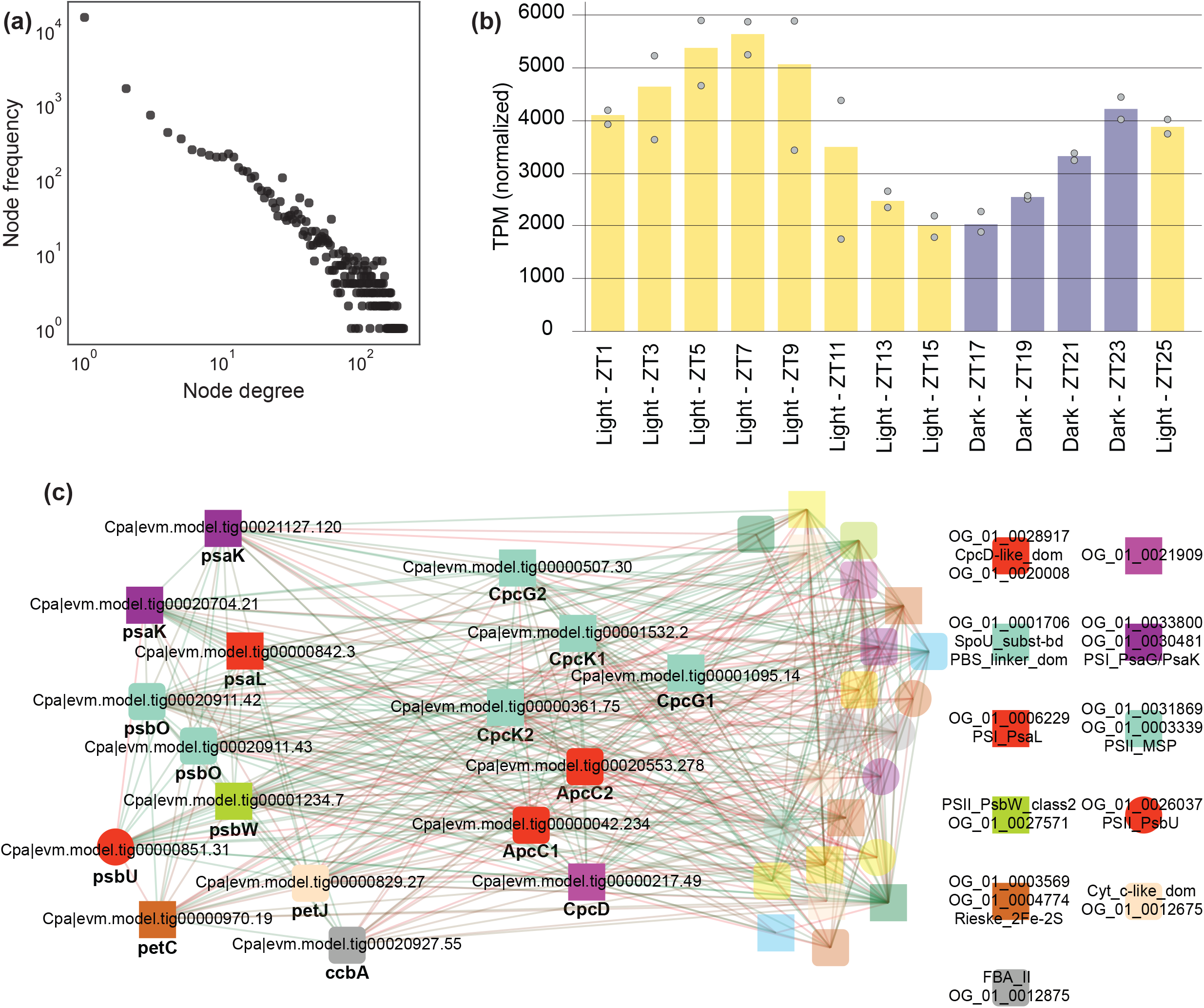
Co-expression network of *Cyanophora paradoxa*. (a) Power law plot based on the Cyanophora gene expression data. The x-axis indicates the node degree (number of connections a given gene has), while the y-axis shows the frequency of a given degree. Both axes were log_10_-transformed. (b) Expression profile of phycobilisome linker protein CpcK2 (*Cpa|evm.model.tig00000361.75*). The x-axis shows the zeitgeber time (hours after the light was switched on), the bars indicate the average expression at a given time, the dots show the maximum and minimum expression of each sample, while the y-axis shows the expression, as transcripts per million (TPM). The color of the bars shows the samples taken in light (yellow) and dark (blue). (c) Co-expression network of CpcK2 (turquoise square). Nodes represent genes, edges connect co-expressed genes (green, orange, red correspond to strong, medium, weak co-expression), while node colors/shapes indicate the gene family and Pfam domains that the genes have in common. Consequently, the genes with the same colored shapes belong to the same family and have the same Pfam domains.

In order to easily browse the expression profiles of genes under different stress conditions, to analyze the Cyanophora co-expression network (Table S10) and functional gene modules, and to compare the gene modules to higher species of the Archaeplastida kingdom, we added our gene expression data, together with publicly available diurnal samples for Cyanophora (Ferrari *et al.*, 2019) to the CoNekT-Plants platform (Proost & Mutwil, 2018). CoNekT-Plants is a user-friendly web-tool containing functional and expression data for 8 species including a glaucophyte (the newly added *Cyanophora paradoxa*), chlorophyte (*Chlamydomonas reinhardtii*), a gymnosperm (*Picea abies*), two monocots (*Oryza sativa*, *Zea mays*) and three dicots (*Vitis vinifera*, *Arabidopsis thaliana* and *Solanum lycopersicum*), and allows comparative genomic and transcriptomic network analysis for these species. CoNekT-plants allows the users to view expression profiles and co-expression networks of their genes of interest, but also performs more sophisticated analyses, such as identification of functional gene modules involved in a biological process of interest, comparison of gene expression across different species, identification of genes highly expressed in a given stress or organ, and others (Proost & Mutwil, 2018).

To exemplify the applicability of the tool for Cyanophora, we explored the expression profile and co-expression network of CpcK2 (this was done by entering *Cpa|evm.model.tig00000361.75* gene identifier to the search box), a gene encoding a linker protein of phycobilisomes (https://conekt.sbs.ntu.edu.sg/sequence/view/4475). *Cyanophora paradoxa* as a glaucophyte lacks the light-harvesting chlorophyll antenna complex present in green algae and higher plants but, similarly to cyanobacteria and red algae, possesses a phycobilin-based light-harvesting antenna complex known as phycobilisomes (PBS) (Watanabe *et al.*, 2012). Phycobilisomes consist of phycobiliproteins associated with linker proteins, which capture and transfer light energy to photosystems I and II (PSI and PSII) (Chang *et al.*, 2015). The analysis of the expression profile of CpcK2 (*Cpa|evm.model.tig00000361.75*) (Fig. 2b) showed an oscillatory pattern with high expression during the light period and a decreased expression in the dark. Interestingly, we also observed an anticipatory behavior (Whitehead *et al.*, 2009), where the expression of the gene decreased towards the end of the day (anticipating the night) and increased towards the end of the night (anticipating the day). The anticipatory behavior is typical for genes under circadian control and is existing to optimize the expression and activity of genes during the day/night cycle (Bell-Pedersen *et al.*, 2005).

We explored the co-expression network associated with the CpcK2 (by clicking on the graph icon under the ‘Co-expression networks/Neighborhood’ panel), which showed genes directly co-expressed (direct neighbors) with the query gene. In the network we identified additional linker protein encoded genes (CpcK1, CpcG1, CpcG2, CpcD, ApcC1, ApcC2) (Watanabe *et al.*, 2012), together with several genes encoding for proteins associated with PSII (psbO, psbW, psbU) (Nickelsen & Rengstl, 2013), PSI (psaK, psaL) (Fromme *et al.*, 2001), cytochrome b6/f complex (petC) (Hasan *et al.*, 2013) and the photosynthetic electron transport (petJ, Fig. 2c) (Durán *et al.*, 2004). Since co-expressed genes are functionally related, the other genes in the network are prime candidates for being involved in photosynthesis. Thus, the database allows to rapidly identify functionally related genes in Cyanophora.

### Functional analysis of network clusters

Functionally related genes form highly connected clusters in the co-expression network, and these clusters can be identified and studied to unravel the functional gene modules of an organism (Mutwil *et al.*, 2010; Rhee & Mutwil, 2014; Aoki *et al.*, 2015) To identify the functional gene modules of Cyanophora, we used the Heuristic Cluster Chiseling Algorithm (HCCA) (Mutwil *et al.*, 2010), and obtained 319 clusters of co-expressed genes (Table S9). To reveal their function, we performed an enrichment analysis and identified 114 clusters significantly enriched for at least one biological process (Fig. 3a, Fig. S3, BH-adjusted empirical p-value < 0.05), as defined by MapMan bin (Schwacke *et al.*, 2019). All the bins were enriched in at least one cluster, except Polyamine metabolism. Other bins such as Redox homeostasis, Chromatin organization, DNA damage response, Protein degradation, Protein translocation, Nutrient uptake, Multi-process regulation, Carbohydrate metabolism, and Nucleotide metabolism were enriched in multiple clusters, suggesting a more complex regulation of these processes involving several gene modules (Fig. S3).

**Figure 3.**
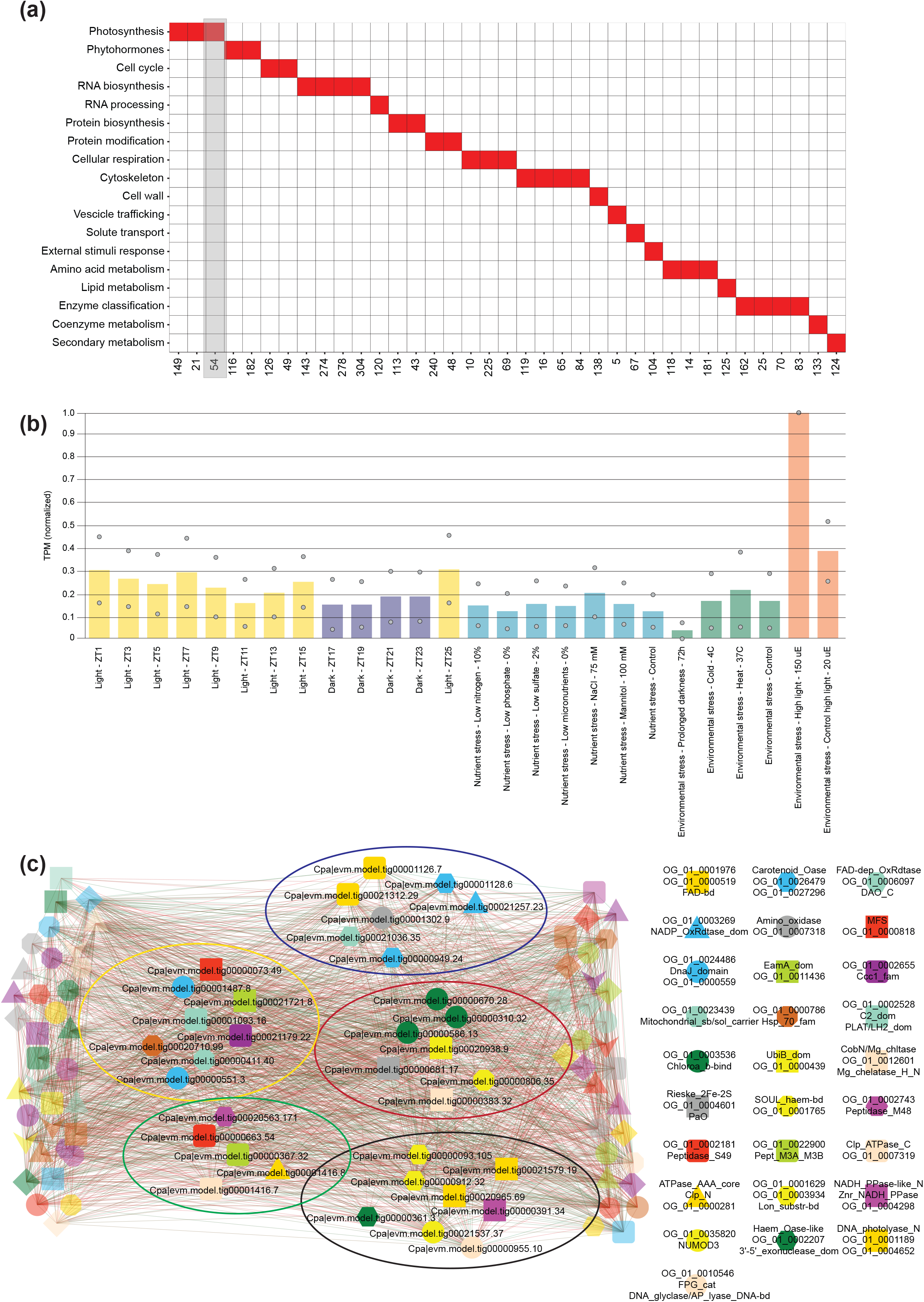
Functional analysis of co-expression clusters of *Cyanophora paradoxa*. (a) The columns represent cluster IDs, while the rows indicate biological processes, as defined by MapMan bins. Clusters enriched for a MapMan bin (FDR adjusted p-value < 0.05), are indicated by red squares. For brevity, only 79 clusters enriched at most for one bin are shown in this figure. (b) Average expression profile of the genes found in cluster 54. The different experiments are indicated on the x-axis and are color-coded according to the type of the experiment, where, e.g., nutrient stress samples are colored light blue. The gene expression levels are indicated on the y-axis. (c) Co-expression network of cluster 54. The genes involved in photosynthesis (red ellipse), oxidoreductases (blue ellipse), peptidases (green ellipse), DNA repair (black ellipse) and transporters (yellow ellipse) are indicated.

CoNekT-Plants shows the average expression of all genes belonging to a given cluster. As an example, we explored cluster 54 (https://conekt.sbs.ntu.edu.sg/cluster/view/45), which was significantly enriched for the process “Photosynthesis” (Fig. 3a, indicated by a gray rectangle), and observed that the cluster expression is the highest for the sample “Environmental stress - High light - 150 μE”. In contrast, “Environmental stress - Prolonged darkness - 72h” showed the lowest expression in this cluster, supporting the evidence of association with photosynthesis-related processes (Fig. 3b). This observation suggests that cluster 54 contains genes that are highly responsive to the amount of light perceived by Cyanophora.

We further explored the network associated with cluster 54 and by looking at the labels obtained from InterPro annotation, we identified five major classes of genes: oxidoreductases, transporters, photosynthesis machinery related proteins, peptidases, and DNA/RNA repair and modification proteins (Fig. 3c). Additionally, Gene Ontology enrichment found on the cluster page for cluster 54 (GO enrichment is found under the expression profile), showed a significant enrichment of terms related to oxidoreduction (“oxidation-reduction process” GO:0055114, “oxidoreductase activity” GO:0016491, “oxidoreductase activity, acting on single donors with incorporation of molecular oxygen” GO:0016701). These results suggest that this light responsive cluster is likely to be active during high light stress and is involved in the cellular repair of biological components damaged by high light. Thus, this example underlines the applicability of co-expression analyses on uncovering functional gene modules.

### Comparative co-expression analysis reveals conserved cell division program

For another case study illustrating comparative co-expression analyses in Cyanophora we selected the cell cycle and its regulation, which have been extensively studied in many model organisms. Notably, the many components involved in DNA replication and chromosome separation are highly conserved across clades and kingdoms of life (Stuart *et al.*, 2003; Yu *et al.*, 2003). However, given the complexity of multicellular organisms, certain aspects of cell division have evolved to accommodate the different morphologies and lifestyles of single and multicellular plants. For example, the coupling of cell division with chloroplast division is found in many unicellular algae but is absent in multicellular plants (Miyagishima *et al.*, 2012). Furthermore, while cell division tends to be regulated by light/dark cycle in algae, multicellular land plants are unlikely to show this behavior (Ferrari *et al.*, 2019)

To study the conservation of cell division in Cyanophora in relation to the plant kingdom, we examined gene *Cpa|evm.model.tig00020554.77* from *C. paradoxa* (https://conekt.sbs.ntu.edu.sg/sequence/view/13961), which is annotated as “DNA replication licensing factor MCM2” and shows the typical cyclic expression of cell division genes, peaking towards the end of the light period during a diurnal cycle (Fig. S4) (Ferrari *et al.*, 2019). MCM proteins contain a Mini-Chromosome Maintenance (MCM) domain and are well known to be involved in DNA replication during both initiation and elongation (Bell & Dutta, 2002). Phylogenetic and expression analysis of the orthogroup of the MCM2 gene (https://conekt.sbs.ntu.edu.sg/tree/view/5112) revealed that the gene has orthologous genes in *C. reinhardtii*, S. moellendorfii *P. abies*, *O. sativa*, *Z. mays*, *V. vinifera*, *A. thaliana*, and *S. lycopersicum*, and that these orthologs are expressed in various tissues in land plants (Fig. 4a).

**Figure 4.**
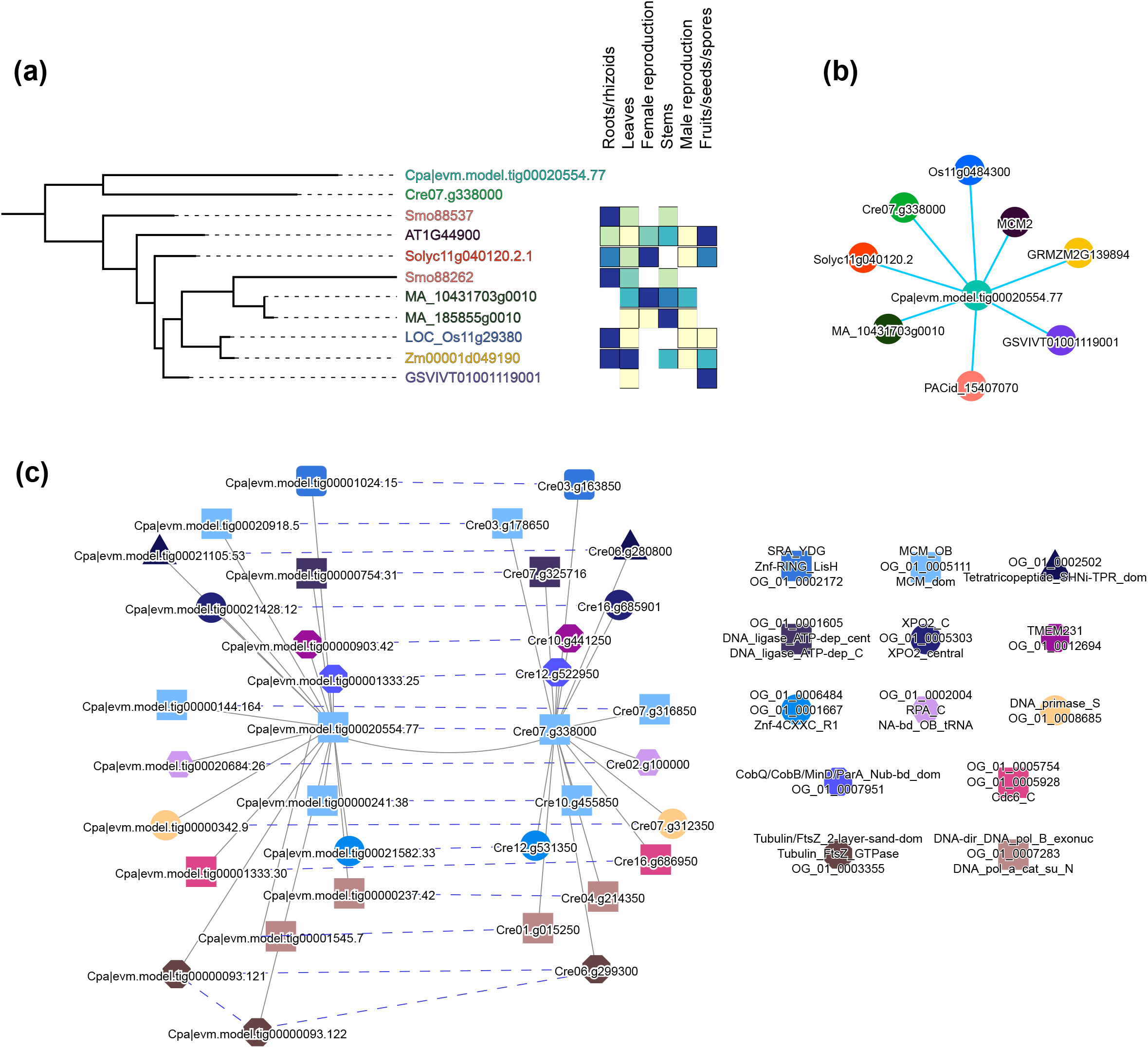
Comparative analysis of cell division gene modules in the plant kingdom. (a) Phylogenetic tree of the gene family OG0005111, which *Cpa|evm.model.tig00020554.77* MCM2 gene belongs to. The genes from different species are color-coded, where, e.g. tomato and rice genes are indicated in red and blue, respectively. The expression of these genes in the various tissues (e.g., roots, leaves, stems) of multicellular plants are shown to the right, where the heatmap indicates high (blue) and low (yellow) expression, respectively. (b) Expression context conservation network of the *Cyanophora* MCM2 gene. Nodes represent gene modules, while similar gene modules are connected by blue edges. The gene modules from the different species are color-coded with the same colors as the genes in the phylogenetic tree. (c) Co-expression networks of the MCM2 modules from Cyanophora (left) and Chlamydomonas (right). Nodes represent genes, solid edges connect co-expressed genes, while dashed edges connect genes belonging to the same family.

CoNekT uses the Expression Context Conservation (ECC) and label co-occurrence values to identify conserved co-expressed gene modules (Movahedi *et al.*, 2011; Proost & Mutwil, 2018). The ECC network showed that the *C. paradoxa* cell division module is conserved across the Archaeplastida kingdom (Fig. 4b, https://conekt.sbs.ntu.edu.sg/ecc/graph/13961/3/1). We then further explored the conserved corresponding module in *C. reinhardtii*, the species which showed the highest ECC value (0.11) to the Cyanophora MCM2 gene. The ECC pair network (https://conekt.sbs.ntu.edu.sg/ecc/graph_pair/759355) showed the presence of several orthologs (dashed lines) and among them, multiple genes with a MCM domain (light blue square). Several other genes with domains related to DNA replication and cell division are found in the network, for example DNA ligase (Moriyama & Sato, 2014), SRA_YDG (Citterio *et al.*, 2004), DNA primase (Moriyama & Sato, 2014), Cdc6 (Castellano *et al.*, 2001), tubulins (Cross & Umen, 2015), DNA polymerase and MinD (Adams *et al.*, 2003; Miyagishima *et al.*, 2012). A closer inspection of the genes present in the network revealed the presence of two orthologs for DNA polymerase A (*Cpa|evm.model.tig00000237.42*, *Cpa|evm.model.tig00000342.9*) and one for DNA polymerase D (*Cpa|evm.model.tig00001545.7*) (Zones *et al.*, 2015) and four orthologs of the MCM protein necessary to form the heterohexamer active during DNA replication (*Cpa|evm.model.tig00020554.77*, MCM2; *Cpa|evm.model.tig00000144.164*, MCM4; *Cpa|evm.model.tig00020918.5*, MCM6; *Cpa|evm.model.tig00000241.38*, MCM7) (Tuteja *et al.*, 2011).

Taken together, these results show how the core components of the cell division machinery are conserved over a great evolutionary distance and showcase how the expression data and the updated CoNekT-plants can be mined to study the conservation of biological processes.

### Comparative analysis of stress responses in algae and flowering plants

Comparative transcriptomic studies across species have helped us considerably to understand which biological pathways are conserved and how they have evolved over time (Mutwil *et al.*, 2011; Movahedi *et al.*, 2012; Hansen *et al.*, 2014; Ruprecht *et al.*, 2017a). For example, we could show that diurnal transcriptional programs are conserved over a huge time span of 1 billion years (Ferrari *et al.*, 2019). The co-expression networks are also conserved across several species (Ruprecht *et al.*, 2011, 2016, 2017a; Mutwil *et al.*, 2011). In the present study we aimed to investigate if the same is true for transcriptomic responses to a wide range of abiotic stresses. To this aim, we retrieved publicly available gene expression data capturing similar stresses as studied in *C. paradoxa* (Table S3) for *Synechocystis sp. PCC 6803*, *C. reinhardtii* and *A. thaliana*. Gene expression data for high light, N deprivation and micronutrient stress is available for all 4 species. Data for sulphate deprivation is published for all species except for *C. reinhardtii*, and dark, heat, cold, phosphate deficiency and salt stress is available for *C. paradoxa* and *A. thaliana.* We also included our data on mannitol stress for *C. paradoxa*.

We first investigated how many genes change their expression profile upon exposure to different stresses. The fraction of stress-specific DEGs is overall below 20% across species, except for prolonged darkness, which results in about 26% and 31% of DEGs in Arabidopsis and Cyanophora, respectively (Fig. 5a, gray bars, Table S4-S7). Other stresses that caused strong transcriptional responses were nitrogen deprivation and temperature stresses. Interestingly, Synechocystis shows an overall large transcriptomic response with many DEGs found for most stresses, suggesting that cyanobacteria might respond more strongly to stresses. We further observed that in most cases the proportion of upregulated and downregulated genes was equal (Fig. 5b, blue/red bars). Few exceptions included light stress in *C. paradoxa* and phosphate and iron deprivation in *A. thaliana* showing a high proportion of upregulated genes, and sulphate deprivation in Synechocystis and salt stress in *A. thaliana* which lead to predominantly downregulated genes.

**Figure 5.**
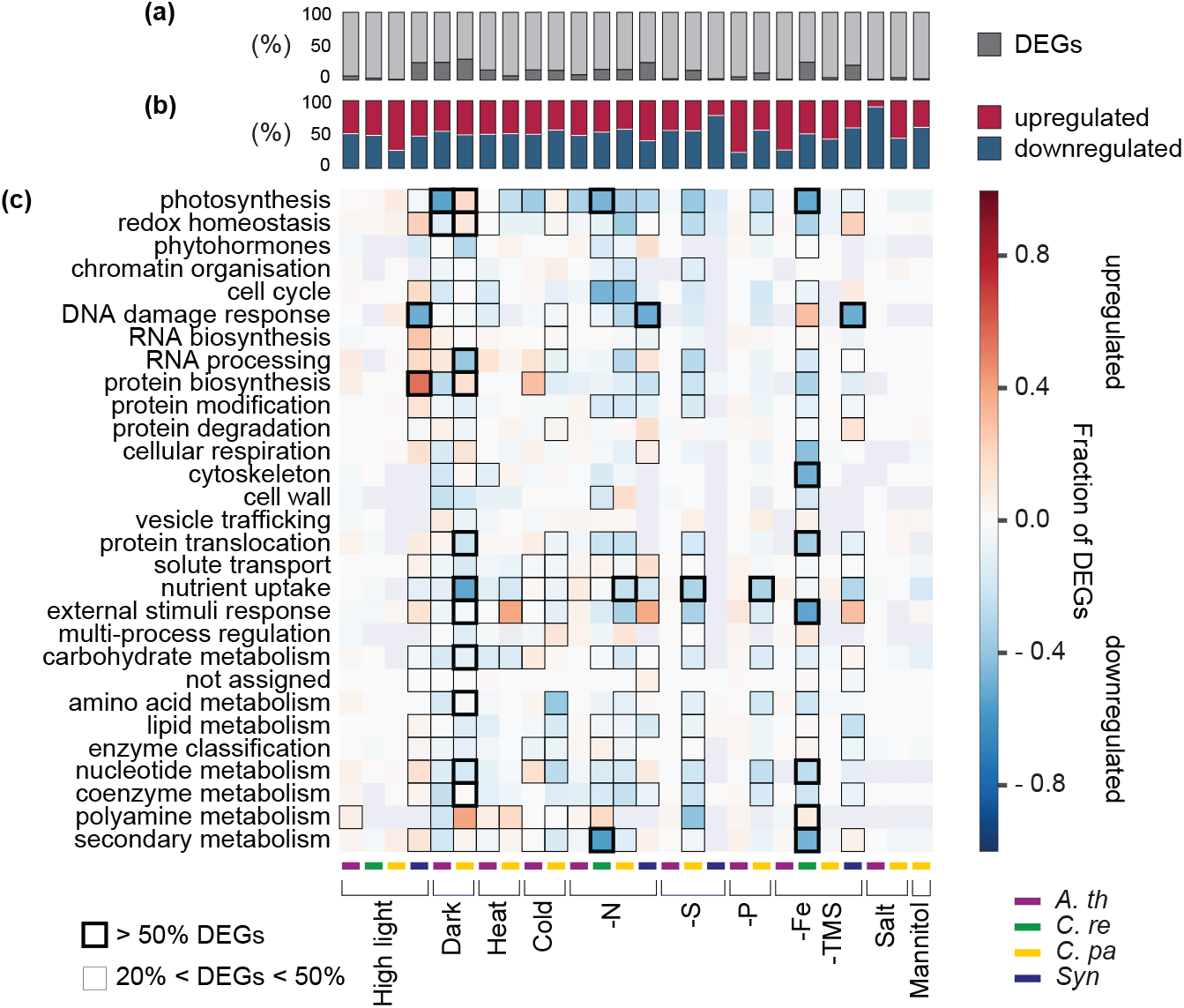
Expression of biological processes upon stress treatment. A) The dark gray bars correspond to the percentage of differentially expressed genes (DEGs) found in the different stresses. B) The red and blue bars indicate the percentage of the upregulated and downregulated genes, respectively. C) The heatmap shows the fraction of differentially expressed genes that are up-(red) or down-regulated (blue), in the different stresses (columns) and biological processes (rows). Thick, thin and no frames surrounding the cells indicate ≥50%, 50%>DEG≥20% and <20% of DEGs in a given biological process/stress combination. The purple, green, yellow and blue bars indicate the species, as shown in the legend.

We then explored how the different biological processes were affected by these stresses. To test that, we analyzed the fraction of DEGs in each of the MapMan ontology bins (Fig. 5c). Prolonged darkness had the strongest impact on the transcriptome, especially for *C. paradoxa*, where more than one third of all biological processes (11 out of 29) showed more than 50% of DEGs (Fig. 5c). The majority of these processes shows a mild prevalence of downregulated genes, suggesting a response to carbon starvation. Interestingly, we observed a general tendency of downregulation of genes involved in photosynthesis for all four species and across all stress conditions.

To test whether stress responses are conserved across species, we measured the similarity of responses of gene families in the different experiments. As a measure of similarity we determined whether a pair of stresses contains more common up- or down-regulated gene families than expected by chance. We found that the down-regulative responses (blue cells) are more frequent than the up-regulative responses (red cells, Fig. 6a). The high light treatment shows very little significant similarity to the other stresses, especially in *C. reinhardtii* and *C. paradoxa*. Conversely, the other conditions showed a significantly similar down-regulatory responses across the stress types and species. For example, the nitrogen depletion response is similar across the four analyzed species, but also similar to the other nutrient depletion conditions, heat, and cold responses (Fig. 6a, blue cells). Overall we conclude that the down-regulatory responses are highly conserved, regardless of the type of stress and species.

**Figure 6.**
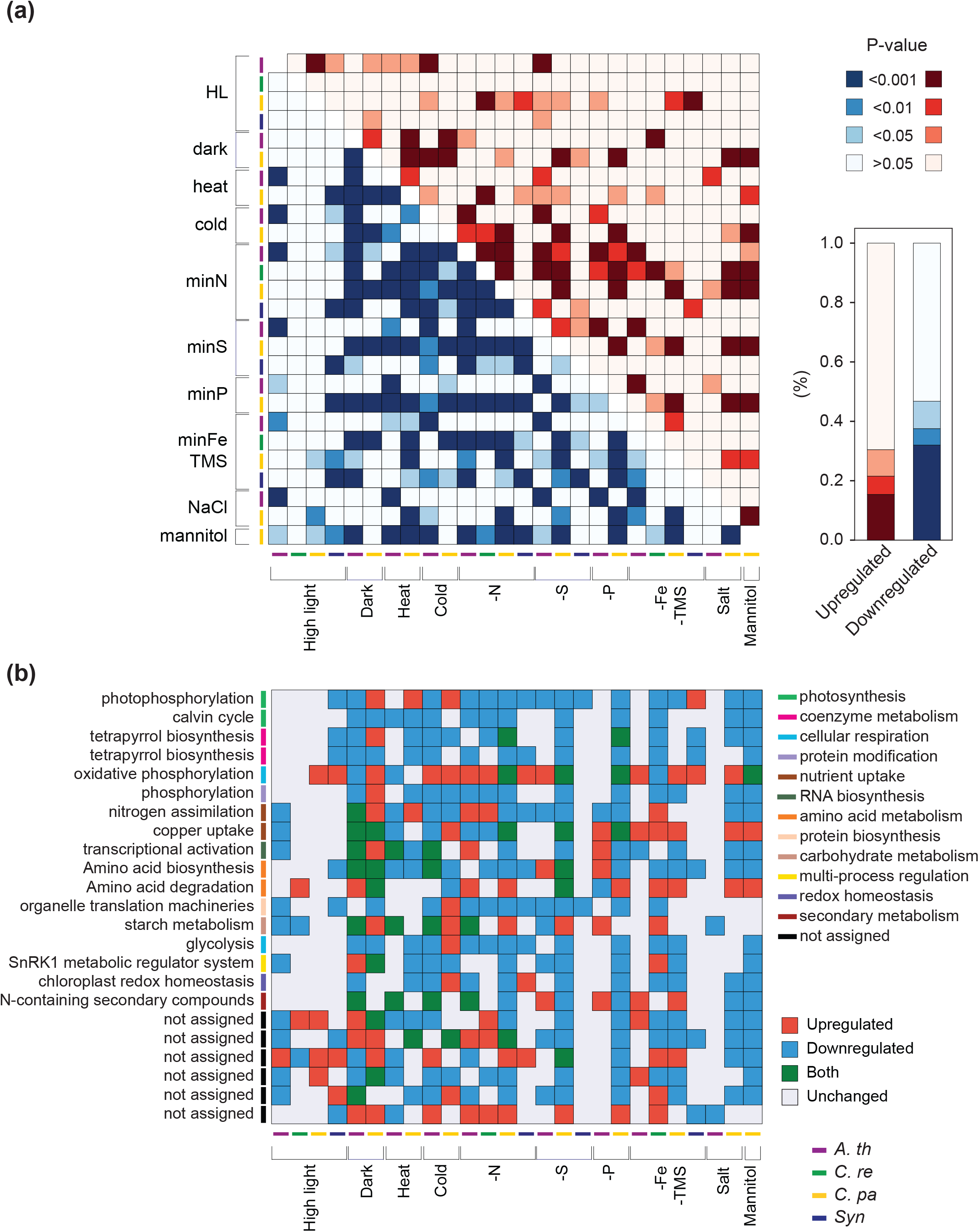
The similarity of the transcriptomic responses to stresses. (a) Conservation of transcriptomic responses of the different stresses and species. The rows and columns indicate the stresses (e.g., dark, heat, cold) and species (e.g., Arabidopsis: purple, Cyanophora: yellow), while the lower left and upper right part of the heatmap indicate the conservation of down- and up-regulated responses, respectively. The intensity of the cell colors indicates the FDR-corrected p-value of conserved responses. (b) Transcriptomic responses of the gene families that are significantly responding to the stresses included in this study. Rows indicate the function of the gene families and columns represent stresses and species. The color of the cells indicate whether the genes in a given family are upregulated (red), downregulated (blue), up- and down-regulated (green) or unchanged (gray) in given stress.

Next, we set to investigate which gene families tend to significantly respond to the stresses (Figure 6b). We observed that gene families involved in photosynthesis, tetrapyrrole biosynthesis, and glycolysis, showed a predominant downregulation response. Other processes, such as nutrient uptake, cellular respiration, amino acid, and protein biosynthesis and modification showed a mixed response to the stresses. Taken together, these results show that the conserved responses across the Archaeplastida kingdom involve downregulation of numerous transcripts, especially photosynthesis, during stress.

### Phylostratigraphic analysis of stress responses

It has been previously shown that the age (phylostrata) of a gene family (Domazet-Loso *et al.*, 2007) is correlated with the expression levels of the corresponding genes in the plant kingdom (Ruprecht *et al.*, 2017a; Ferrari *et al.*, 2019) and the number of publications dedicated to the gene family (Ruprecht *et al.*, 2017a; Hansen *et al.*, 2018). The latter finding suggests that older gene families produce more pronounced mutant phenotypes and are involved in more fundamental biological processes. It is unclear, however, whether gene families that appeared at a specific time in plant evolution are more stress responsive.

To investigate whether the age of a gene family is correlated to its stress response, we analyzed the fraction of genes that is either up- or downregulated in each phylostratum (i.e., the relative age of the gene family). We arranged the phylostrata from oldest (prokaryotic) to youngest (species-specific gene families) and observed that the response to stress is evenly distributed across all phylostrata for both up- and down-regulated genes. Among them, the oldest phylostratum (prokaryotic) showed the highest enrichment for the downregulation response, being significantly enriched in >50% of the experiments. Furthermore, the Gymnospermae phylostratum, representing gene families that appeared in the ancestor of seed plants, tends to be upregulated in heat, salt, low iron, and phosphate. Conversely, species-specific genes showed the least enrichment, suggesting that newly formed genes are less involved in the stress response. We conclude that stress responses tend to be regulated independently from the age of the gene families, except the oldest (prokaryotic, more responsive than expected) and the youngest (species-specific, less responsive than expected).

## Discussion

*Cyanophora paradoxa* is the only glaucophyte representative with a sequenced genome, and due to the importance of this species for evolutionary studies we generated a comprehensive expression atlas comprising several abiotic stresses. We focused the atlas on a wide range of abiotic stresses for two reasons. The first is that co-expression networks require gene expression data capturing as many different observations as possible (Usadel *et al.*, 2009). The second is that all organisms have developed transcriptional programs to cope with various abiotic stresses (Zhu, 2016), but currently no study is available on gene expression comparison at plant kingdom-wide scale.

The Cyanophora gene expression atlas revealed that there was no correlation between the severity of growth inhibition caused by the stress and the number of transcriptional changes (given as the number of differentially expressed genes) (r=0.278, p-value=0.437, Fig. 1e), even though both culture growth and the transcriptome were affected by the applied stresses. For instance, stresses that cause a mild growth retardation (e.g. nitrogen and sulphate deprivation) showed a high number of DEGs, while stresses causing a severe growth inhibition (salt and mannitol) showed the lowest amount of DEGs. The overall low correlation between growth inhibition and transcriptome changes suggests highly diverse strategies and regulatory programs that control the stress acclimation.

We expanded the CoNekT-Plants database with the newly generated expression dataset to give easy access to study gene expression and co-expression networks through the use of various comparative tools. These tools allow us to overcome the paucity of knowledge related to gene function in *C. paradoxa*. We exemplify these tools by studying phycobilisome formation (Fig. 2c) and high light response (Fig. 3c), demonstrating how the co-expression network can be used to predict the function of unknown genes. Furthermore, the updated CoNekT-Plants can be used for cross-species analyses, as we have demonstrated by identifying a conserved module involved in cell division in *C. paradoxa* and *C. reinhardtii* (Fig. 4c). These tools thus allow to uncover the similarities and differences in gene expression and functional modules across evolutionary distances, making CoNekT-Plants a unique resource to study the evolution of functional modules.

Moreover, we analyzed whether the responses to abiotic stresses are conserved across conditions and organisms, and we observed a conserved downregulation response related to genes involved in photosynthesis and primary metabolism (Fig. 5c, 6). This observation is also supported by the analysis of the most conserved families operating upon stress (Fig. 6b), in which we find several, mostly downregulated photosynthesis and primary metabolism-related families. Similar trends have been found in grasses, where the various members of the Pooideae family showed highly diverse transcriptomic responses to cold stress, but a common downregulation of transcripts involved in photosynthesis and metabolism (Schubert *et al.*, 2019). Analysis of photosynthesis parameters during salt stress revealed that this is reflected by a decrease of the total performance index of photosynthesis in multiple species (Pavlović *et al.*, 2019), showing a correlation between the transcript responses and plant phenotype. Our findings suggest that photosynthesis is downregulated to prevent photoinhibition and cellular damage not only in cold and salt stress but for a wider variety of stresses. Furthermore, since this response is seen in algae and land plants, we propose that downregulation of photosynthesis and metabolism is a kingdom-wide, stress-responsive program.

In contrast to the conserved downregulation responses, we observed only a modest conservation of upregulation responses to various stresses across different species (Fig. 6a). This is also reported for salt stress in six Lotus accessions, where only 1% of genes showed a conserved response (Sanchez *et al.*, 2011), in two strawberry cultivars, which displayed a modest conservation of DEGs to the same pathogen (Wang *et al.*, 2017), in Poaceae, which revealed a poor expression conservation of orthologs across same tissue types (Davidson *et al.*, 2012), and in seven Arabidopsis accessions which showed a divergent response of genes to exogenous salicylic acid (van Leeuwen *et al.*, 2007). We propose two non-exclusive hypotheses for this surprising lack of conservation in transcriptomic responses. First, assuming that orthologous sets of genes are still needed for acclimation to a given stress, investigating only one layer of the gene activity regulation (transcript levels) might be insufficient. Second, a high rate of rewiring, divergence, and the presence of multiple signal transduction pathways might confound the signal. To provide a deeper understanding of stress responses, we suggest that snapshots of additional regulatory layers, preferably closer to the end-product of gene regulation (i.e., the protein), should be included in future studies.

Finally, we demonstrated that the oldest gene families, in contrast to the youngest species-specific families, are actively responding to abiotic stress (Fig. 7). This suggests that the young gene families are not typically involved in stress acclimation, which is in line with the fact that the investigated stresses, and the corresponding coping mechanisms, are ancient.

**Figure 7.**
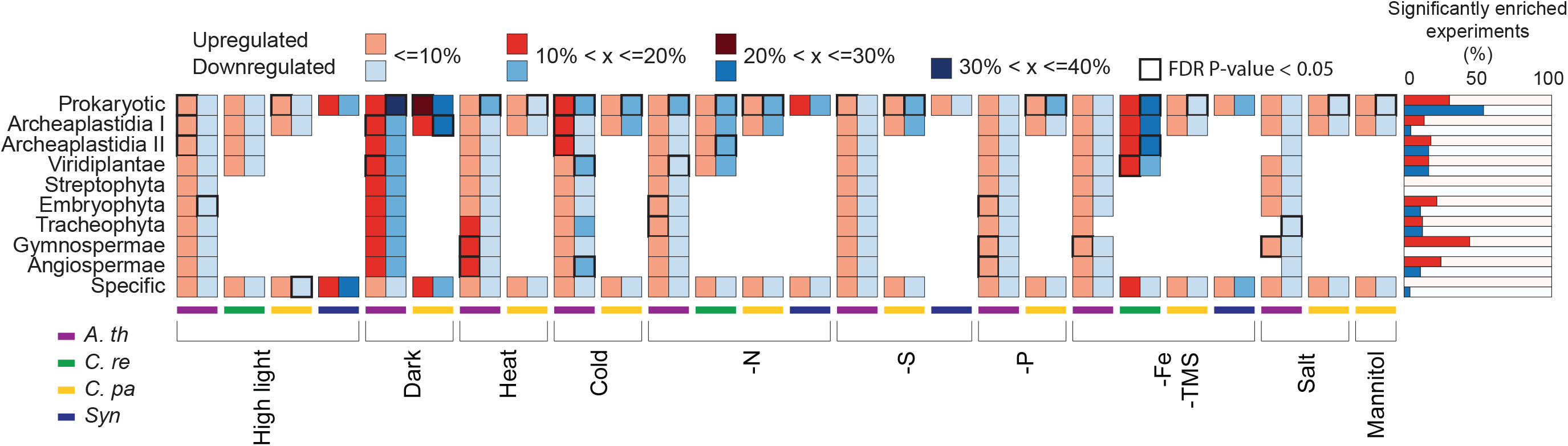
Phylostratigraphic analysis of stress responses. The species and stresses are shown in columns, while the phylostrata, arranged from oldest (prokaryotic) to youngest (species-specific) are shown in rows. The colors indicate the percentage of genes that are up- (red) or down-regulated (blue) in a given phylostratum. The black cells indicate phylostrata that are significantly enriched in given stress and species.

Thus, our study on abiotic stress responses in *C. paradoxa* provides new insights on stress responses, gene function prediction, and conservation and evolution of abiotic stress acclimation.

## Acknowledgments

We would like to thank Dr. Daniela Mutwil-Anderwald for proofreading this article.

## Author contributions

M.M. conceived the project. C.F. and M.M. designed the study. C.F. performed the experiments and data analysis with input from M.M. C.F. and M.M. wrote the manuscript.

---

**Figure S1. Quantification of starch in prolonged darkness.** Grey and yellow bars represent average optical density (OD, 680nm) measurements for three independent samples exposed to prolonged darkness (72h) and continuous light, respectively. Error bars indicate the standard deviation.

**Figure S2. PCA analysis of *C. paradoxa*.** Each dot represents an RNA-seq sample, and the different colors represent a growth condition for (a) nutrient stresses and (b) environmental stresses.

**Figure S3. Significantly enriched clusters of *C. paradoxa*.** The columns represent cluster IDs, while the rows indicate biological processes, as defined by MapMan bins. Clusters enriched for a MapMan bin (FDR adjusted p-value < 0.05), are indicated by red squares.

**Figure S4. Diurnal expression profile of MCM2 (*Cpa|evm.model.tig00020554.77*).** The x-axis shows the zeitgeber time (hours after the light was switched on), the bars indicate the average expression at a given sample, the dots show the maximum and minimum expression of each sample, while the y-axis shows the expression, as transcripts per million (TPM). The color of the bars shows the samples were taken in light (yellow) and dark (blue).

**Figure S5. Commonly enriched families across abiotic stresses and species.** Transcriptomic responses of the gene families that are significantly (FDR corrected empirical p-value < 0.05) responding to the stresses included in this study. Rows indicate the gene families and columns represent stresses and species. The color of the cells indicate whether the genes in a given family are upregulated (red), downregulated (blue), up- and down-regulated (green) or unchanged (gray) in given stress.

**Table S1. Composition of the C media.**

**Table S2. LSTrAP mapping statistics.** The first column represents the sample names, the second, third and fourth columns represent the reads mapped to the genome, the number of reads mapped to noncoding genes (‘no feature’) and not uniquely (‘ambiguous’), respectively. The fifth, sixth, and seventh columns indicate the relative percentages. The eighth column indicates the percentage of reads mapped to coding genes.

**Table S3. Gene expression data used in this study.** The table indicates the species, stress, experiment identifier, platform type, and stress description.

**Table S4-S7. A table containing up- and downregulated genes in all stresses and species.** The first column shows the gene name, the second column indicates the condition the gene resulted differentially expressed, the third and fourth columns indicate the adjusted p-value and the log2 fold change associated to the gene and the experiment.

**Table S8. *Cyanophora paradoxa* TPM-normalized gene expression matrix.**

**Table S9. HCCA clusters of *C. paradoxa***

**Table S10. Co-expression network of *C. paradoxa***

## Notes

https://conekt.sbs.ntu.edu.sg/

