## Supplementary figures and images for "Gene expression analysis of *Cyanophora paradoxa* reveals conserved abiotic stress responses between basal algae and flowering plants"

### Fig S1

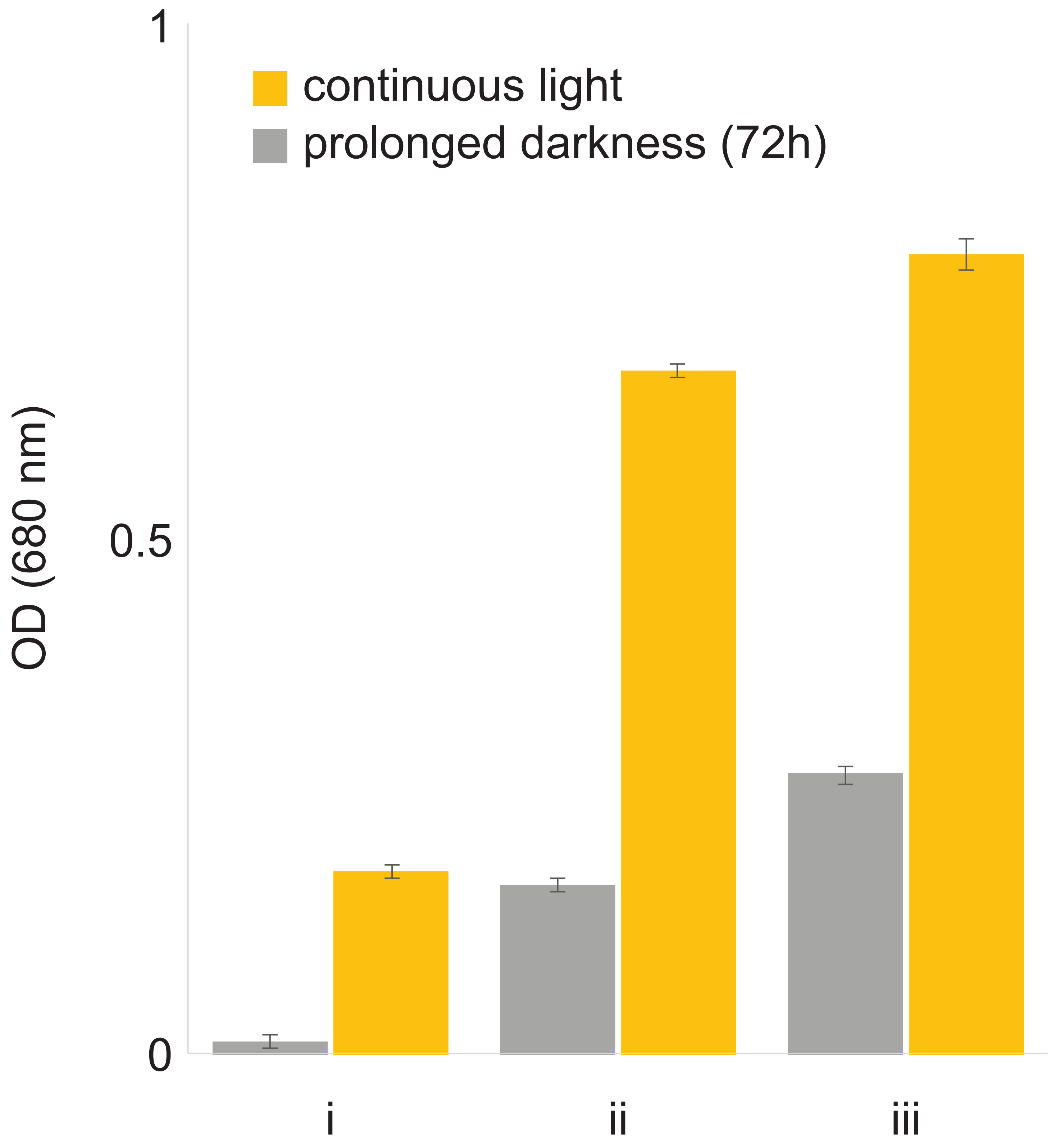

### Fig S2

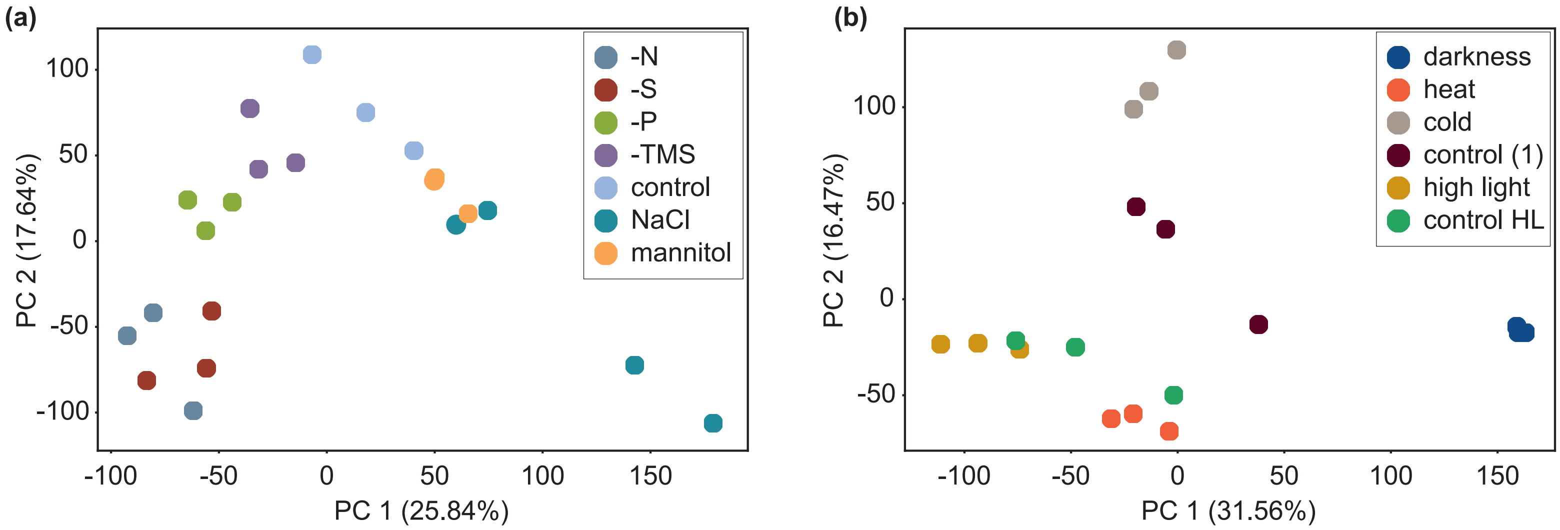

### Fig S3

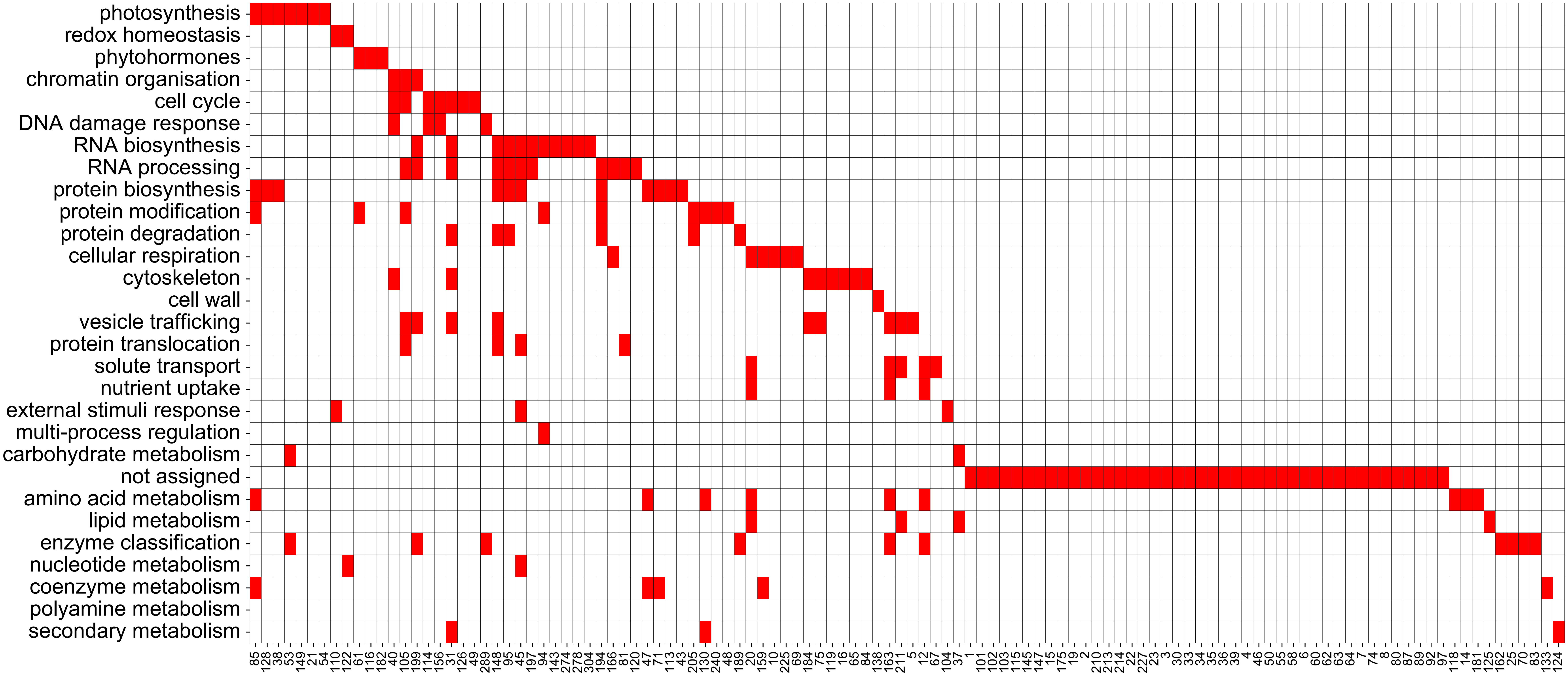

### Fig S4

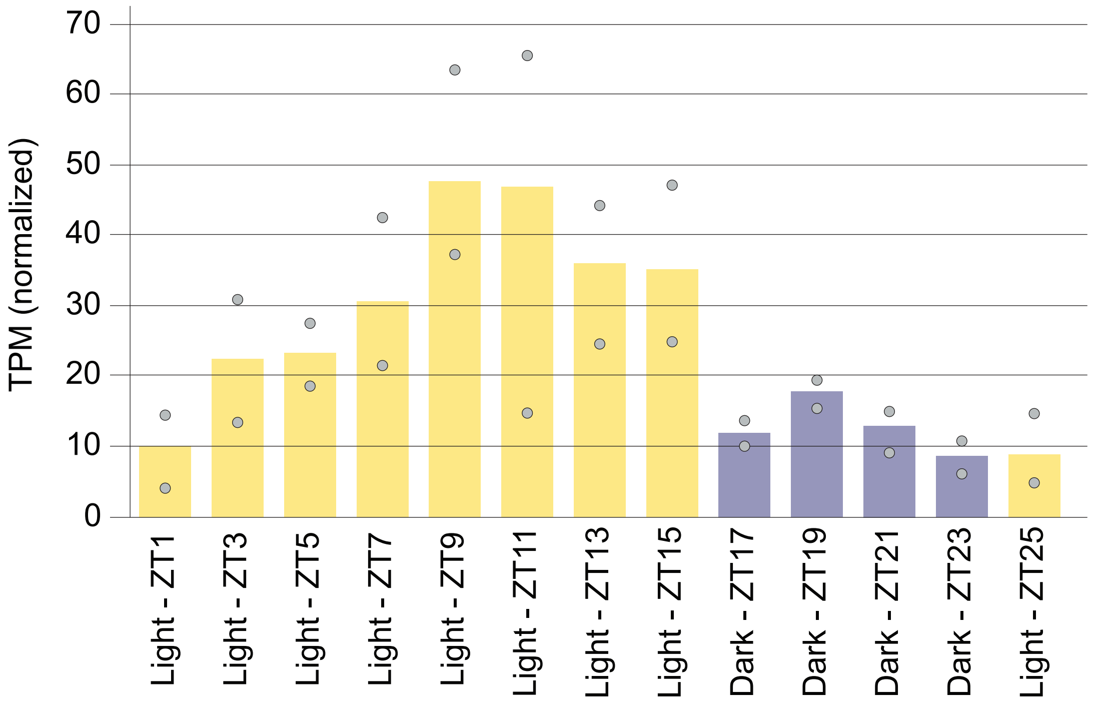

### Fig S5

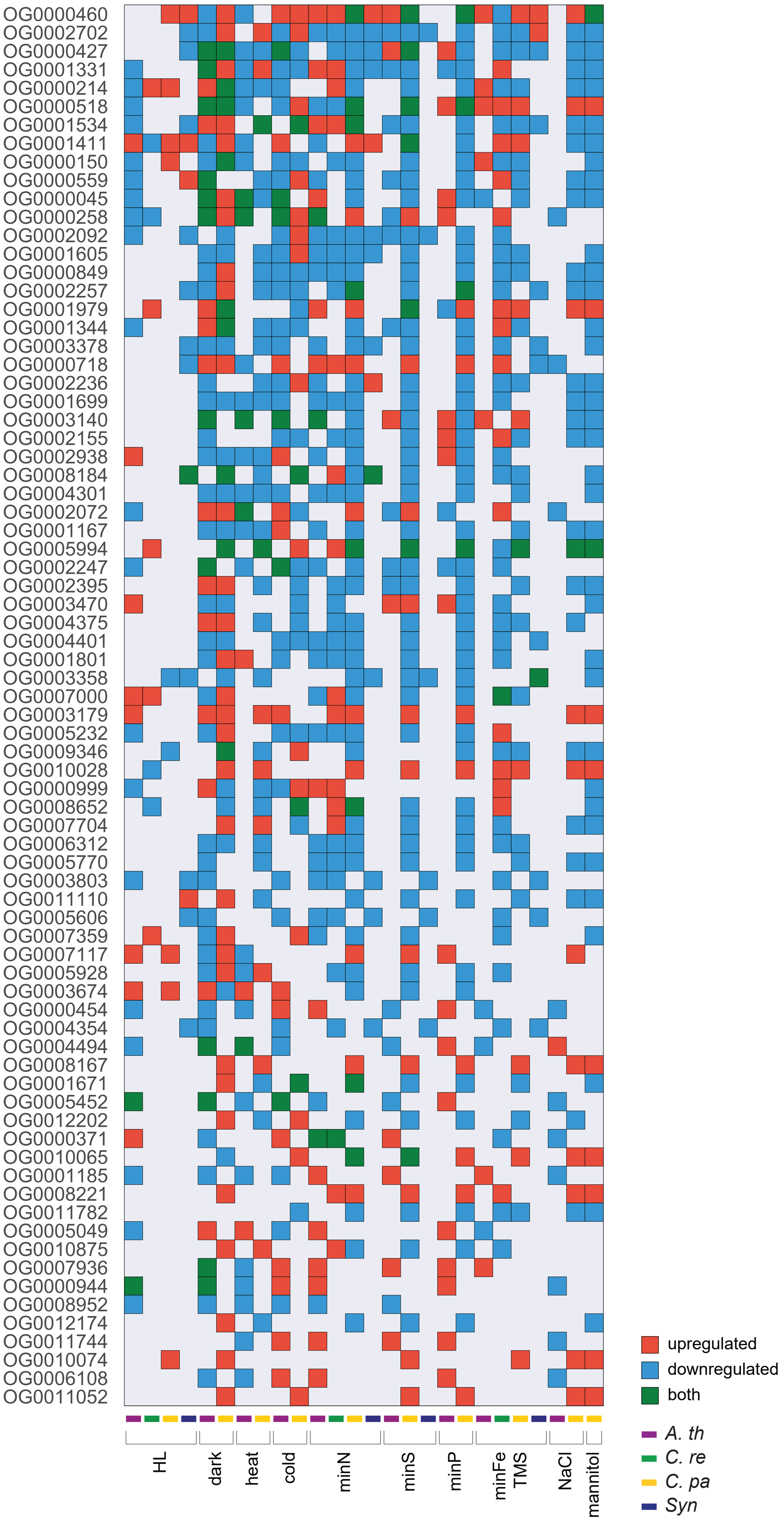
